# Intracellular Bismuth Coordination of Peptides and Miniproteins

**DOI:** 10.64898/2026.01.08.698318

**Authors:** Andrew Brennan, Jody M Mason

**Author notes:** Address correspondence to JMM or AB (;).

## Abstract

Cyclisation of peptides has emerged as a powerful methodology to enhance molecular stability and functional efficacy. Bismuth(III) coordination represents a particularly compact and chemoselective cyclisation modality, yet to date has remained confined to *in vitro* or phage-based systems. Here we report the first example of bismuth-mediated peptide and minprotein cyclisation occurring within living cells, achieved directly during recombinant expression in *E. coli*. Supplementation of growth media with bismuth salts enables efficient intracellular coordination of either three or six cysteine residues, yielding bicyclic and tetracyclic architectures with minimal cellular toxicity. Bis-bismuth coordination generated a tetra-cyclic minprotein with exceptional thermal and serum stability, representing a level of intracellular structural reinforcement not previously accessible. This chemistry enables the intracellular constraint of peptides and proteins and is directly compatible with genetically encoded peptide and miniprotein libraries, establishing bismuth coordination as a compact and robust modality for live-cell molecular screening.

## Introduction

Peptides and miniproteins have emerged as promising therapeutic modalities capable of engaging protein-protein interaction surfaces that are often challenging for small molecules.^1-3^ Cyclisation has become a widely adopted strategy to enhance peptide efficacy, improving target engagement, biostability and in some cases cellular uptake.^4-6^ In parallel, chemical modification approaches have expanded the range of peptide topologies accessible in drug discovery, enabling exploration of conformationally restricted architectures. Cyclisation strategies have facilitated access to macrocyclic and bicyclic peptides during screening campaigns, contributing to the identification of several *de novo* candidates that have progressed toward clinical evaluation.^7, 8^ A variety of chemistries have been developed to enable peptide cyclisation, including ring closing metathesis, lactamisation and cysteine-directed alkylation reactions.^9-11^ More recently, bismuth mediated cyclisation has been shown to generate peptide bicycles under mild conditions, with applications in the development of cell-penetrating peptides and in cyclic peptide phage display libraries.^12-14^ Bismuth is particularly attractive in this context due to due to its strong thiophilicity, enabling rapid and chemoselective coordination to cysteine residues, and its ability to act as a compact single-atom cyclisation scaffold. Peptide bicycles constrained by bismuth coordination have been reported to exhibit enhanced stability, including prolonged persistence in buffered solution and reduced susceptibility to proteolytic degradation compared to linear peptides.

Genetically encoded peptide library screening provides a powerful means to explore vast sequence spaces and identify molecules with desirable biological activities.^15, 16^ Integrating chemical modifications or cyclisation strategies into such libraries further expands their functional diversity and increases the likelihood of discovering potent and selective candidates.^17, 18^ We have developed the Transcription Block Survival (TBS) peptide library screening platform, which utilises live bacterial cells to enable competition selection of peptides that antagonise protein-DNA interactions.^19-21^ Building on this approach, we have shown that certain cysteine-selective bis-alkylating reagents can be supplemented into bacterial growth media, enter live cells, and cyclise recombinantly expressed peptides and proteins, facilitating intracellular cyclisation-coupled TBS (icTBS) screening. Using this strategy we previously identified a sidechain-to-sidechain cyclised peptide antagonist of the oncogenic transcription factor CREB1 from a 63-million-member library, with demonstrated activity in in cancer cell models.

Here, we show that *E. coli* cells can tolerate optimised concentrations of bismuth salts, enabling bismuth entry into the cytoplasm and chemoselective coordination with cysteine residues in recombinantly expressed peptides and proteins. By extending this chemistry to miniprotein scaffolds, we demonstrate that bismuth coordination can enhance structural stability, yielding constrained architectures compatible with intracellular functional assays, including icTBS. Together, these findings establish intracellular bismuth coordination as a versatile strategy for constraining peptides and proteins in living cells and provides a foundation for the development of genetically encoded peptide and minprotein libraries with therapeutic potential.

## Results and discussion

To investigate intracellular bismuth cyclisation we first generated a test peptide containing three cysteines separated by diverse amino acid spacers to validate chemoselectivity, expressed with an N-terminal 6xHis-SUMO tag to facilitate expression and purification (**Figure 1A**). Notably, no additional cysteine residues are present in the conjugated construct. Two bismuth(III) salts previously shown to cyclise cysteine-containing peptides *in vitro* (BiBr_3_ and BiK_3_[citrate]_2_) were evaluated.^12, 14^ Bacterial growth assays were performed to identify bismuth concentrations compatible with recombinant expression and intracellular cyclisation (**Figure 1B-1E**). While BiBr_3_ exhibited higher toxicity than BiK_3_[citrate]_2_, *E. coli* cells were able to grow in the presence of both compounds under appropriate conditions.

**Figure 1.**
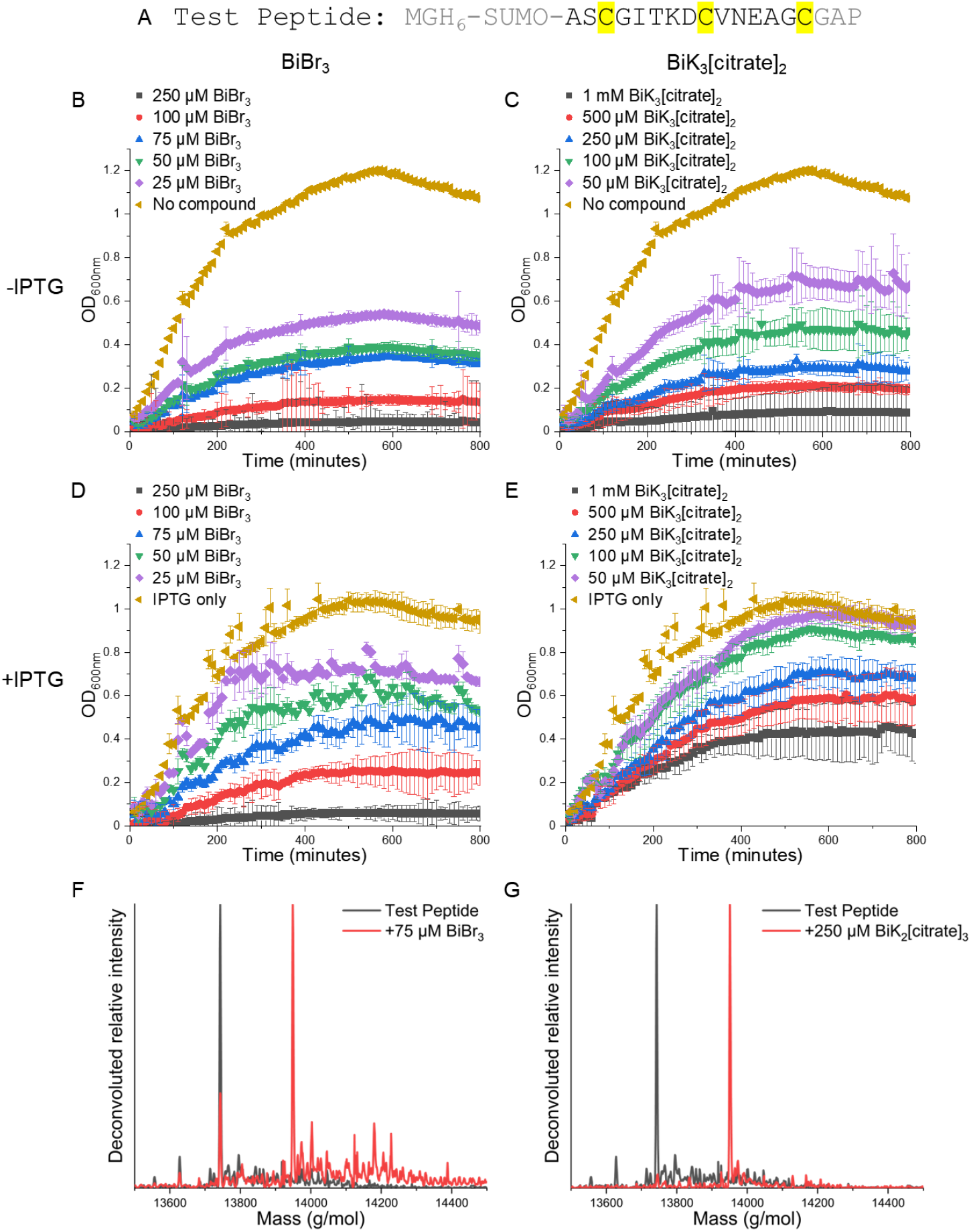
Intracellular bismuth-mediated cyclisation of a model peptide in E.coli. (A) Sequence of the Test Peptide expressed as an N-terminal His_6_–SUMO fusion for intracellular cyclisation (B-E) E. coli growth curves (OD_600_) measured during incubation with increasing concentrations of bismuth salts, illustrating cellular tolerance to both compounds and reduced toxicity for BiK_3_[citrate]_2_ relative to BiBr_3_. (F, G) ESI-MS spectra of purified SUMO-tagged Test Peptide following recombinant expression in the presence of bismuth salts, showing mass shifts consistent with coordination of a single bismuth atom. N-terminal methionine truncation was observed upon recombinant expression.^22^

To assess intracellular cyclisation under these conditions, bacterial cultures expressing the 6xHis-SUMO-Test Peptide were grown in the presence of bismuth salts across a range of concentrations, with IPTG added to induce recombinant expression. Peptides were purified by immobilised metal affinity chromatography and analysed by LC-MS to determine cyclisation efficiency. Cyclisation was quantified using the cyclic-to-linear peptide ratio derived from MS peak intensities as off-target reactions were not observed but cannot be excluded. For both compounds, peptide cyclisation was detected at all concentrations tested, with mass shifts corresponding to coordination of a single bismuth atom (**Figure 1F, 1G, Table S1**). As cell pellets were washed prior to lysis to remove unreacted bismuth salt, all detected cyclised peptide must have been generated within living bacterial cells.

We evaluated intracellular cyclisation as a function of compound concentration, balancing cyclisation efficiency against bacterial growth. BiBr_3_ exhibited higher toxicity and a narrower usable concentration range, resulting in lower achievable cyclic-to-linear peptide ratios, whereas BiK_3_[citrate]_2_ was better tolerated and supported substantially higher cyclisation efficiencies across the concentrations tested (Figure 1F,G; Table S1). On this basis, 75 µM BiBr_3_ (cyclic-to-linear ratio of 3, indicating a reaction yield of ∼75%) and 250 µM BiK_3_[citrate]_2_ (cyclic-to-linear ratio of 21, indicating a reaction yield of ∼96%) were identified as the highest concentrations compatible with robust bacterial growth and efficient intracellular cyclisation and were taken forward for further analysis.

To assess whether efficient cyclisation was maintained throughout the growth window relevant to intracellular screening, cultures expressing the test peptide were supplemented with 75 µM BiBr_3_ or 250 µM BiK_3_[citrate]_2_, with samples taken at ∼0.1 OD_600nm_ intervals (Figure 2). In both cases, high cyclic-to-linear peptide ratios were observed throughout growth. Consistent with previous intracellular cyclisation studies, cyclisation efficiency decreased gradually during growth, reflecting depletion of compound from the media. However, robust levels of cyclised peptide were retained at the at the growth endpoint corresponding to icTBS passaging, with 75 µM BiBr_3_ producing a cyclic-to-linear ratio of 10.9 and 250 µM BiK_3_[citrate]_2_ producing a ratio of 26.0 (indicating reaction yields of ∼92% and ∼96% respectively). Taken together, these data establish 250 µM BiK_3_[citrate]_2_ as the optimal condition for generating bismuth-constrained peptides and proteins within live *E. coli*.

**Figure 2.**
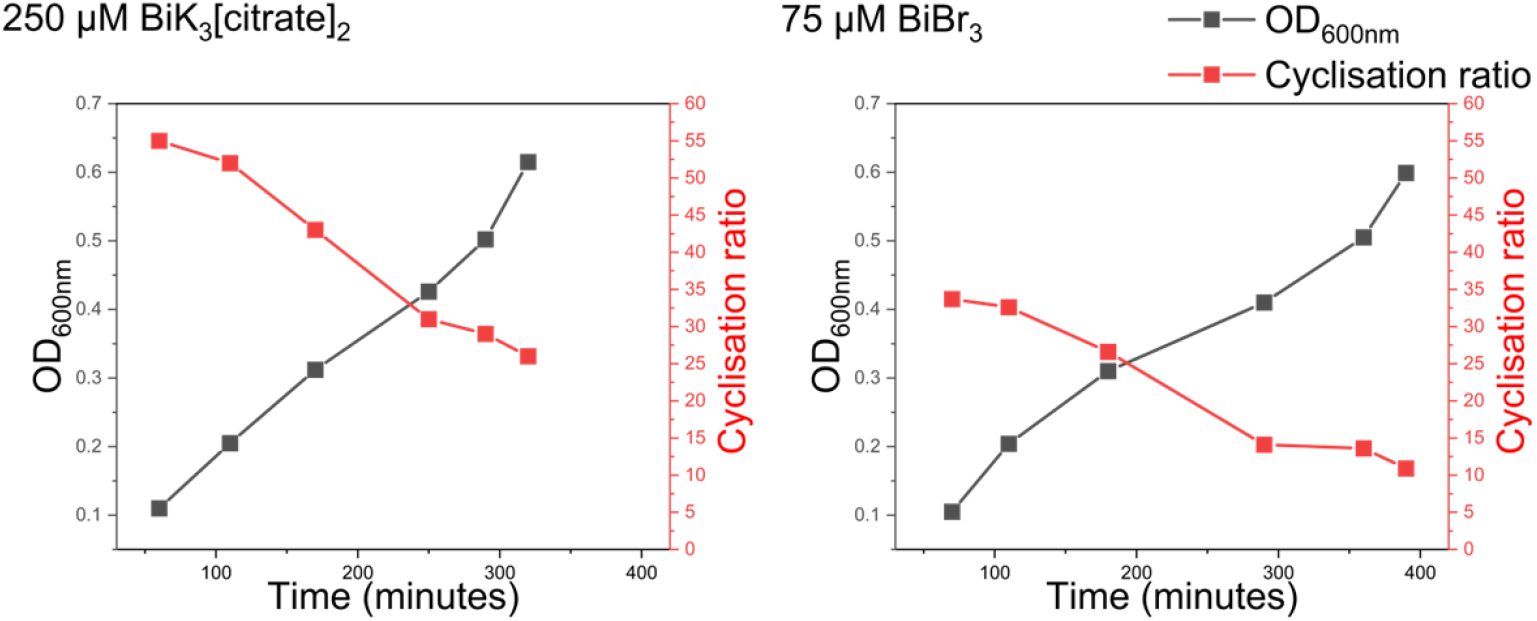
Intracellular bismuth-mediated peptide cyclisation is maintained throughout bacterial growth. Intracellular cyclisation efficiency of the Test Peptide during E. coli growth in the presence of 250 µM BiK_3_[citrate]_2_ or 75 µM BiBr_3_. Cultures were sampled throughout growth to assess both bacterial density (OD_600_) and cyclic-to-linear peptide ratios by LC–MS. Under both conditions, efficient peptide cyclisation was maintained across the growth window relevant to intracellular screening.

To expand the topological complexity of constrained constructs, we next explored the use of bismuth coordination to chemically constrain miniproteins, with the aim of enabling intracellular screening of structurally diverse libraries using icTBS. We distinguish peptides and miniproteins by their ability to adopt stable 3D tertiary structures, with miniproteins typically forming compact, folded cores, whereas peptides generally remain more conformationally flexible in solution. Guided by a series of computationally designed disulphide-constrained miniprotein reported by Bhardwaj *et al*..^23^, we selected a three helix bundle containing two disulphide bonds (gHHH_06, PDB ID: 2ND2) as a scaffold for modification. From this parent structure, three miniprotein variants were designed to introduce either one or two cysteine triads to enable bismuth coordination (**Figure 3**). For **Miniprotein1**, one cysteine pair from gHHH_06 was removed and an additional cysteine was added in proximity to the remaining cysteine pair, to introduce a cysteine triad for cyclisation. **Miniprotein2** design removed both cysteine pairs from the original design and a new triad was introduced with two cysteines introduced at *i→i+4* positions in one helix and a third cysteine in the neighbouring helix. For **Miniprotein3** two triads were introduced by adding additional cysteines in proximity to the cysteine pairs in the original design, with C3 in the gHHH_06 sequence moved to the second position to accommodate the bismuth atom within the structure. Where cysteines were removed from the original design they were replaced with alanine residues. Alphafold predictions indicated that all designs retained the intended three-helix fold, with minimal perturbation relative to the parent scaffold.^24^

**Figure 3.**
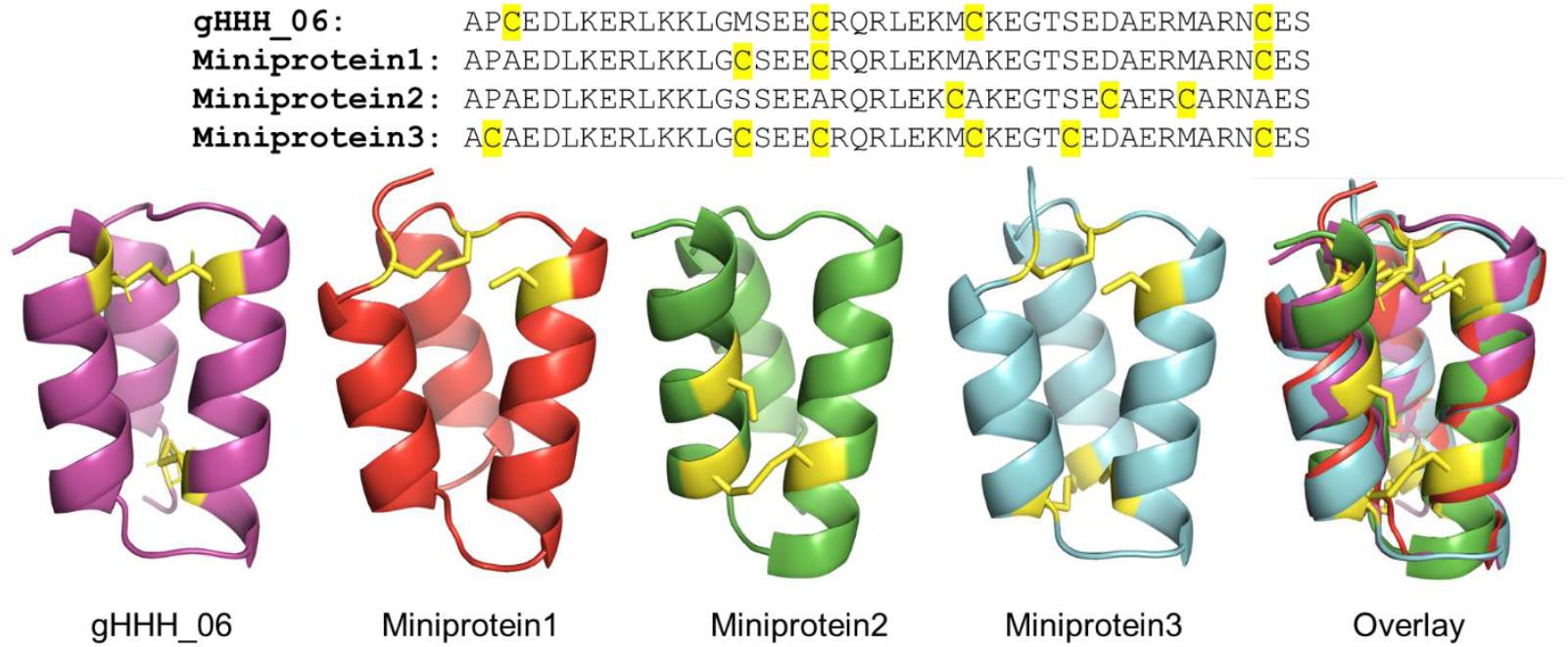
Design and structural prediction of bismuth-coordinated miniproteins. Sequences and predicted structures of designed miniproteins based on a three-helix bundle scaffold, illustrating cysteine placements intended to enable bismuth coordination. The parent gHHH_06 structure reported by Bhardwaj et al. is shown for reference (PDB ID: 2ND2). Miniprotein structures are AlphaFold predictions, with predicted three-helix bundle topologies retained across all designs (PTM scores: 0.69, 0.61, and 0.74 for Miniproteins 1, 2, and 3, respectively).^24^

The secondary structure of synthesised miniproteins (**Figure S1-3**) was assessed by circular dichroism (CD) spectroscopy in both linear and bismuth-constrained forms, in the presence or absence of the reducing agent TCEP (**Figure 4A, Figures S1-S4**). All three bismuth-constrained miniproteins exhibited increased alpha-helicity, with enhanced minima at 222 nm. **Miniprotein3** Cyclised displayed the greatest helicity, with characteristic minima at 208 and 222 nm and a fractional helicity of 62% relative to 29% in the linear form. Addition of TCEP had no effect on the bismuth-constrained miniproteins, consistent previously reported bismuth-constrained peptide resistance to glutathione reduction.^13^ TCEP reduced helicity for the linear constructs, indicative of the formation of stabilising disulphides in the absence of bismuth.

**Figure 4.**
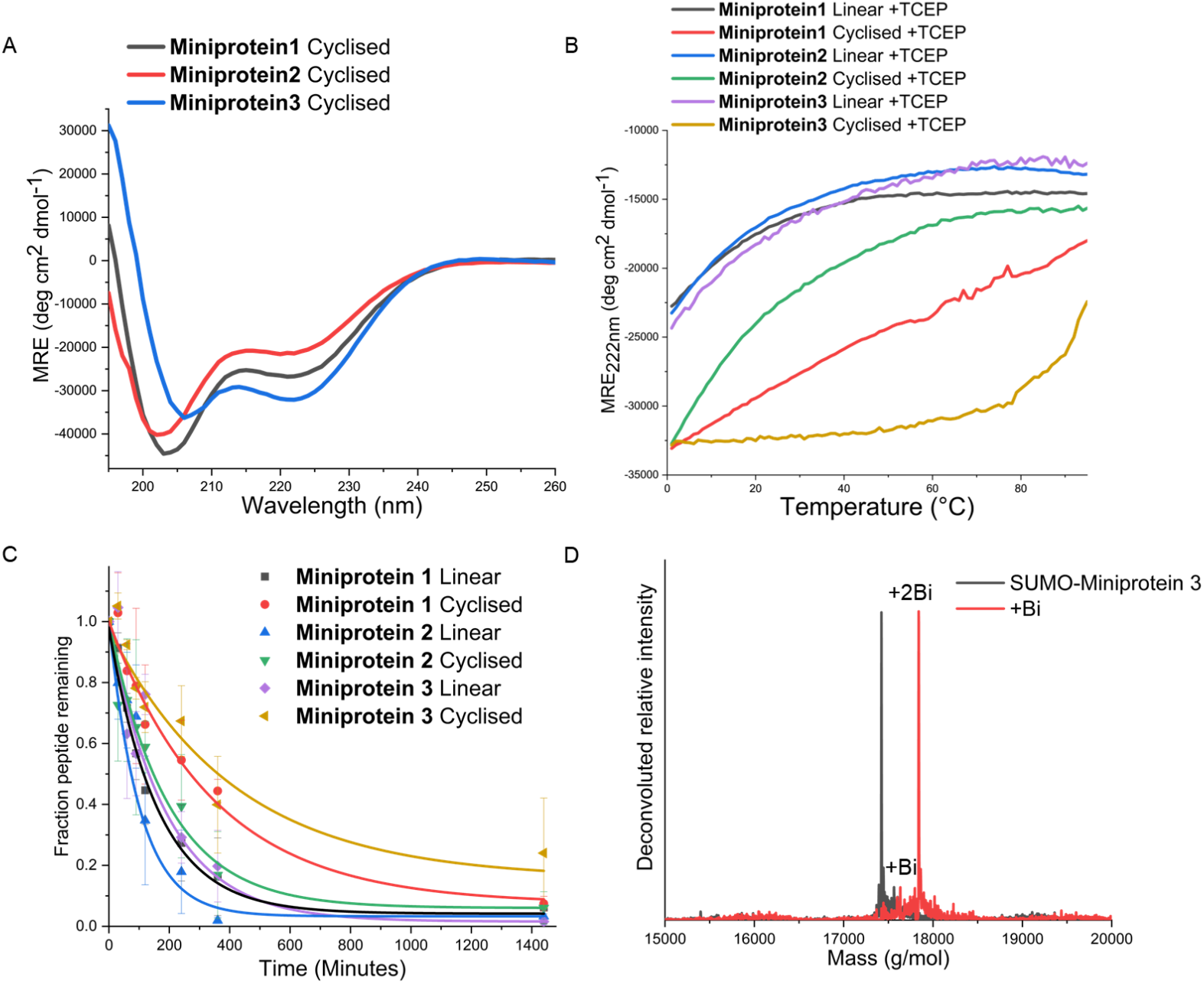
Biophysical characterisation and intracellular generation of bismuth-constrained miniproteins. (A) CD spectra of bismuth-constrained miniproteins (100 µM), showing increased α-helicity relative to linear counterparts, with Miniprotein3 Cyclised exhibiting the highest helical content. (B) Thermal denaturation CD profiles demonstrating enhanced thermal stability upon bismuth coordination, with Miniprotein3 being most pronounced. Linear miniproteins display similar melting behaviour, indicating minimal perturbation of the underlying fold following sequence modification. (C) Human serum stability of linear and bismuth-constrained miniproteins measured by ESI-MS over 24 h, showing increased persistence upon bismuth coordination, particularly for Miniprotein3. (D) ESI-MS spectra of SUMO-Miniprotein3 following recombinant expression in E. coli in the presence or absence of 250 µM BiK_3_[citrate]_2_, revealing mass shifts consistent with coordination of two bismuth centres and formation of a tetra-cyclic construct during intracellular expression. N-terminal methionine truncation was observed upon recombinant expression.^22^

Thermal denaturation experiments further demonstrated that bismuth coordination enhanced miniprotein stability (**Figure 4B**). For Miniproteins 1 and 3, cyclisation increased thermal stability, with the doubly coordinated **Miniprotein3** retaining secondary structure up to ∼80 °C. While **Miniprotein2** exhibited increased helicity upon cyclisation, no corresponding increase in thermal stability was observed.

To assess biological stability, human serum degradation assays were performed over 24 h (Figure 4C; Figure S5). In agreement with the CD data, bismuth coordination significantly increased serum stability for all three constructs, with doubly constrained **Miniprotein3** showing the greatest persistence (t_1/2_ = 361 ± 28 min).

**Table 1.** Structural and biological stability of designed miniproteins. Fractional α-helicity was determined by circular dichroism spectroscopy at 222 nm under reducing conditions at 37 °C. Serum stability was assessed by incubation in human serum at 37 °C, with intact miniprotein quantified by LC–MS to derive half-life values. Data are reported as mean ± standard deviation (n = 3). Cyclised constructs correspond to bismuth-coordinated miniproteins.

|  | Fraction helicity at 37°C<br>(Reduced) | Serum half-life (minutes) |
| --- | --- | --- |
| **Miniprotein1** Linear | 33% | 115±7 |
| **Miniprotein1** Cyclised | 52% | 283±11 |
| **Miniprotein2** Linear | 29% | 96±16 |
| **Miniprotein2** Cyclised | 41% | 149±12 |
| **Miniprotein3** Linear | 33% | 131±7 |
| **Miniprotein3** Cyclised | 62% | 361±28 |

To confirm that bismuth-cyclised miniproteins can be generated within living cells, plasmid constructs encoding 6xHis-SUMO-tagged Miniproteins1 and 3 were expressed in *E*.*coli* in the presence of 250 µM BiK_3_[citrate]_2_. **Miniprotein2** was not pursued further due to its lower stability. Following purification, ESI-MS analysis of the SUMO-tagged constructs revealed mass increases consistent with coordination of one or two bismuth atoms (Figure 4D, S6). In the case of **Miniprotein3**, this corresponds to dual bismuth coordination, generating a tetra-cyclic construct directly within living cells.

Circular dichroism spectroscopy of cleaved and purified miniproteins demonstrated that recombinantly expressed, intracellularly cyclised constructs adopt secondary structures highly similar to those of the corresponding synthetic proteins (Figures S7 and S8). Together, these results demonstrate that structurally reinforced, bismuth-constrained miniproteins can be generated during recombinant expression in *E. coli*, providing a foundation for the development of genetically encoded miniprotein libraries compatible with intracellular screening platforms such as icTBS.

## Conclusions

We have demonstrated that supplementation of bismuth salts into bacterial growth media enables efficient cyclisation of peptides and miniproteins directly within *E. coli*. This strategy supports the formation of multi-cyclic architectures around compact, single atom coordination scaffolds and extends our previous work on intracellular chemical constraint using bis-alkylating agents. Together, these approaches expand the range of chemically constrained peptide and miniprotein topologies that can be accessed in living cells and applied to intracellular screening platforms.

Miniproteins have shown clear therapeutic potential through their potent target binding affinities, facilitated by expansive interaction surfaces, and increased stability when compared to peptides and antibodies. Computational, grafting and screening approaches have all produced miniproteins capable of effectively engaging with target proteins, highlighted by the FDA approval of the phage display-derived miniprotein Ecallanitide.^25-28^ Here, we show that bismuth coordination increases the chemical, thermal, and serum stability of miniproteins, including the generation of highly stable tetra-cyclic architectures. These findings support the use of chemically constrained miniproteins as robust scaffolds for genetically encoded library design and intracellular functional screening.

More broadly, intracellular bismuth-mediated coordination provides a straightforward and well-tolerated route to constrain peptides and miniproteins during recombinant expression. By enabling chemically reinforced architectures to be generated directly inside living cells, this work expands the scope of intracellular peptide and miniprotein library screening approaches, including those based on Transcription Block Survival. As such, bismuth coordination offers a compact and versatile constraint for the discovery of peptide and protein-based antagonists with translational potential.

## Acknowledgments

JMM is grateful to the Biotechnology and Biological Sciences Research Council (BB/X001849/1, and BB/T018275/1).

