## Supplemental Information for "Intracellular Bismuth Coordination of Peptides and Miniproteins"

### Materials and Methods

**Bacterial growth assays:** These growth assays were performed to establish bismuth salt concentrations compatible with intracellular peptide cyclisation while maintaining sufficient bacterial viability for downstream screening applications. As demonstrated previously for intracellular cyclisation using cysteine-selective bis-alkylating agents, overexpression of cysteine-containing peptides can partially mitigate compound toxicity by acting as intracellular sinks. For this reason, growth was assessed both in the presence and absence of IPTG induction. In the context of intracellular screening platforms such as icTBS, reduced growth rates are acceptable provided cultures remain viable and capable of recombinant expression over the screening window**.** pET21a plasmid for the expression of the Test Peptide was transformed into BL21 (DE3) *E. coli* and stored as a polyclonal glycerol stock. To measure compound toxicity, an overnight culture from this glycerol stock was inoculated into 100 µL M9 minimal media per well in a 96-well plate to a starting $OD_{600}$ of 0.05, in the presence of variable bismuth salt (0-250 µM $BiBr_3$ (Merck) or 0-1 mM $BiK_3[citrate]_2$ (Merck)) and IPTG (0 or 1 mM) concentrations. Growth was monitored by measuring $OD_{600}$ over time in a CLARIOstar microplate reader (BMG).

**Intracellular cyclisation test reactions:** Intracellular cyclisation efficiency was assessed using LC-MS by calculating the ratio of cyclic to linear peptide peak intensities rather than absolute reaction yields. Absolute intracellular yields are difficult to determine due to variable expression levels and recovery efficiencies; however, relative peak intensities provide a robust metric for comparing cyclisation efficiency across conditions. Off-target reactions were not observed by LC-MS but cannot be excluded. Prior to cell lysis, pellets were extensively washed to remove unreacted bismuth salt, ensuring that all detected cyclised peptide was generated within living cells rather than during downstream processing. An overnight culture of the Test Peptide or SUMO-Miniprotein1 or SUMO-Miniprotein3 expressing polyclonal stock was inoculated in 500 mL M9 minimal media with 1 mM IPTG and bismuth salt (0-250 µM $BiBr_3$ or 0-1 mM $BiK_3[citrate]_2$) 37 °C with shaking (200 rpm) for 24 hours. For measurement of cyclisation throughout the growth (**Figure 2**), cells were inoculated in 1 L of media and $OD_{600}$ was monitored, with 100 mL of culture taken as a timepoint sample at ~0.1 $OD_{600}$ intervals. Time-course sampling was performed to confirm that effective intracellular cyclisation was maintained throughout the bacterial growth window relevant to intracellular screening. In icTBS assays, cells are typically passaged at OD600 ≈ 0.6; therefore, sustained cyclisation at this growth endpoint is required to ensure that constrained peptides are present during functional selection. Changes in cyclic-to-linear ratios over time reflect progressive depletion of bismuth salt from the media as expression proceeds. Cells were harvested by centrifugation, washed, and lysed, and His-tagged peptides were partially purified via $Ni^{2+}$-affinity chromatography (IMAC). Peptide-containing fractions were analysed by liquid chromatography–mass spectrometry (LC-MS) to determine cyclisation ratio.

**Miniprotein design rationale:** Miniprotein constructs were derived from the disulphide-constrained three-helix bundle gHHH_06 by rational repositioning or replacement of cysteine residues to enable bismuth coordination while preserving the underlying fold. Where cysteine residues were removed from the parent design, alanine substitutions were introduced to maintain helix propensity and minimise structural perturbation. For Miniprotein 1 and Miniprotein 2, single cysteine triads were introduced to enable trivalent bismuth coordination, whereas Miniprotein 3 was designed to accommodate two such triads. These designs enable bismuth coordination but do not assume a fixed coordination geometry, as the primary objective was to generate structurally reinforced scaffolds compatible with intracellular screening rather than rigidly defined architectures.

**Miniprotein structure prediction:** AlphaFold Server was utilised to predict miniprotein structures, with 5 models generated. AlphaFold predictions were used to assess preservation of the overall fold topology following sequence modification rather than to infer atomic-level structural detail. PTM scores in the range observed here (0.61–0.74) are consistent with confident prediction of global fold architecture for small, designed proteins. The retention of predicted three-helix bundle structures across all designs is supported experimentally by circular dichroism and stability measurements, which together validate the use of these scaffolds for intracellular screening applications. Miniprotein1 models had pairwise RMSD values between 0.479 and 0.787 Å (average RMSD: 0.653 Å) and a PTM score of 0.69. Miniprotein2 models had pairwise RMSD values between 0.906 and 2.144 Å (average RMSD: 1.513 Å) and a PTM score of 0.61. Miniprotein3 models had pairwise

RMSD values between 0.397 and 0.760 Å (average RMSD: 0.604 Å) and a PTM score of 0.74.

**Miniprotein synthesis/purification:** Miniproteins were synthesised using a Liberty Prime microwave peptide synthesiser (CEM) at a 0.1 mmol scale on rink amide ProTide resin using standard Fmoc solid-phase methodology. Coupling was performed in a 5 mL reaction using 5x amino acid, 5x Oxyma Pure and 10x N,N'-diisopropylcarbodiimide in dimethylformamide (DMF). Deprotection was performed by addition of 0.75 mL 30% pyrrolidine into the coupling reaction. N-terminal acetylation was performed with 3x acetic anhydride, 4.5x diisopropylethylamine in DMF for one hour at room temperature. Incubation in a cleavage mixture (82.5% trifluoroacetic acid, 2.5% DODT, 5% anisole, 5% trimethylsilyl chloride, 5% dimethyl sulfide, 5% triisopropylsilane, 10mL) for 4 h at room tempearture cleaved the peptide from the resin and removed side chain protecting groups. The resin was removed by filtration and cleaved peptides were precipitated in diethyl ether at -80°C and centrifuged. Peptide was washed a further four times with diethyl ether before it was dried overnight at room temperature. Peptides were resuspended in 1:1 water:acetonitrile (0.1% TFA) before purification using RP-HPLC with a Luna C18(2) column (5-μm particle size, 100 Å pore size, 250 × 21.2 mm; Phenomenex) using a water:acetonitrile gradient (0.1% TFA). Peptide masses and purity were verified by electrospray ionisation mass spectrometry.

SUMO fusion was employed to enhance soluble expression and recovery of miniprotein constructs during recombinant production. Miniprotein 2 was not pursued further in intracellular expression experiments due to its comparatively lower thermal and serum stability observed in vitro, whereas Miniproteins 1 and 3 exhibited enhanced stability upon bismuth coordination and were therefore selected for validation of intracellular cyclisation. For recombinant production, pET21a plasmid for the expression of SUMO-Miniprotein1 or SUMO-Miniprotein3 was transformed into BL21 (DE3) *E. coli* and stored as a polyclonal glycerol stock. An overnight culture from the glycerol stock was inoculated into 1 L of LB media at 37 °C with shaking (200 rpm) until mid-log phase ($OD_{600}$ ≈ 0.6), at which point expression was induced with 1 mM IPTG. Cultures were incubated for a further 1 hour at 37 °C, followed by the addition of 250 μM $BiK_3[citrate]_2$. Cultures were then incubated at 25 °C overnight with shaking. Cells were harvested by centrifugation, washed, and lysed, and His-tagged SUMO-miniprotein fusions were partially purified via $Ni^{2+}$-affinity chromatography (IMAC). SUMO-tagged miniproteins were buffer exchanged into 20 mM Tris.HCl, 1 mM DTT, pH 8.0. A 15:1 mixture of SUMO-tagged miniprotein:ULP1 was incubated at 30°C for 16h. ULP1 was also purified to ~80% purity as above. As constructs were N-terminally His-tagged on the SUMO and the ULP1 has a 6xHis tag also, these impurities were removed by passing the sample through a $Ni^{2+}$-affinity column. The flowthrough containing cleaved miniprotein was finally purified by RP-HPLC as above.

**Bismuth cyclisation**: Purified miniprotein samples were resuspended at 1 mM in a solution of 10 mM Tris-HCl, 10 mM TCEP, pH 7.5. The sample pH was confirmed and adjusted if necessary due to residual TFA from cleavage, before incubation at room temperature for 30 minutes to ensure disulphide reduction. A 100 mM $BiBr_3$ stock was prepared in DMSO and added to the reduced miniprotein to a final concentration of 1.2 mM. The reaction was carried out at room temperature with gentle agitation (100 rpm) for 10 minutes before centrifugation at 15,000 xg for 10 minutes. The constrained miniproteins were purified from the supernatant by RP-HPLC as above.

**Circular dichroism spectroscopy:** An Applied Photophysics Chirascan was used for CD measurements, with a 200 µL 100 µM sample in a 1 mm path length CD cell. Samples were suspended in 20 mM potassium phosphate,150 mM potassium fluoride, with or without 5 mM TCEP.HCl at pH 7.4. Three scans between 190 and 260 nm were collected with a bandwidth of 1 nm and data sampled at a rate of 0.5 $s^{-1}$. These scans were averaged and converted to molar residue ellipticities (MRE). Thermal denaturation experiments were performed by measuring the ellipticity at 222 nm over a 1 to 95°C gradient at 1°C increments. Post-melt scans at 20°C confirmed the transitions were reversible as they overlaid within 8% of the pre-melt scan. The resistance of bismuth-constrained miniproteins to reduction by TCEP is consistent with previous reports that bismuth-coordinated peptide bicycles remain stable in the presence of excess glutathione. While the intracellular redox environment differs from the conditions used here, these observations collectively support a chelate-driven stabilisation mechanism for bismuth coordination. No single intracellular mechanism is proposed. Instead, the data demonstrate that bismuth-mediated constraints remain intact under both chemical reduction and biologically relevant conditions.

**Serum stability:** Miniprotein stocks (250 µM) were prepared in water and 120 µL was added to 480 µL human serum (Merck) before incubation at 37°C. 50 µL aliquots were removed at designated timepoints and added to 200 µL acetonitrile+1%TFA. Samples were vortexed for 30 seconds, incubated at 4°C for 10 minutes and then centrifuged (15000 xg, 10 minutes). The supernatant was analysed by LC-MS to quantify intact miniprotein. Data were collected in triplicate and are plotted as an average with error bars shown as one standard deviation.

**Intracellular cyclisation plasmid constructs:**

**DNA and protein sequence of Test Peptide construct:**

5’-
ATGGGTCATCACCATCATCATCACGGGTCGGACTCAGAAGTCAATCAAGAAGCCAAGCC
AGAGGTCAAGCCAGAAGTCAAGCCTGAGACTCACATCAATTTAAAGGTGTCCGATGGAT
CTTCAGAGATCTTCTTCAAGATCAAAAAGACCACTCCTTTAAGAAGGCTGATGGAAGCGT
TCGCTAAAAGACAGGGTAAGGAAATGGACTCCTTAAGATTCTTGTACGACGGTATTAGAA
TTCAAGCTGATCAGACCCCTGAAGATTTGGACATGGAGGATAACGATATTATTGAGGCTC
ACCGCGAACAGATTGGAGGTGCTAGCTGCGGCATTACCAAAGATTGCGTGAACGAAGC
GGGCTGCGGCGCGCCTTGA-3’

MGHHHHHHGSDSEVNQEAKPEVKPEVKPETHINLKVSDGSSEIFFKIKKTTPLRRLMEAFAK
RQGKEMDSLRFLYDGIRIQADQTPEDLDMEDNDIIEAHREQIGGASCGITKDCVNEAGCGAP
*

**DNA and protein sequence of SUMO-Miniprotein1 construct:**

5’-
ATGGGTCATCACCATCATCATCACGGGTCGGACTCAGAAGTCAATCAAGAAGCCAAGCC
AGAGGTCAAGCCAGAAGTCAAGCCTGAGACTCACATCAATTTAAAGGTGTCCGATGGAT
CTTCAGAGATCTTCTTCAAGATCAAAAAGACCACTCCTTTAAGAAGGCTGATGGAAGCGT
TCGCTAAAAGACAGGGTAAGGAAATGGACTCCTTAAGATTCTTGTACGACGGTATTAGAA
TTCAAGCTGATCAGACCCCTGAAGATTTGGACATGGAGGATAACGATATTATTGAGGCTC
ACCGCGAACAGATTGGAGGTGCTAGCGCGCCGGCGGAAGATCTGAAAGAACGTCTGAA
AAAACTGGGCTGCAGCGAAGAATGCCGCCAGCGCCTGGAAAAAATGGCCAAAGAAGGC
ACCAGCGAAGATGCGGAACGCATGGCGCGCAACTGCGAAAGCTGA-3’

MGHHHHHHGSDSEVNQEAKPEVKPEVKPETHINLKVSDGSSEIFFKIKKTTPLRRLMEAFAK
RQGKEMDSLRFLYDGIRIQADQTPEDLDMEDNDIIEAHREQIGGASAPAEDLKERLKKLGCS
EECRQRLEKMAKEGTSEDAERMARNCES*

**DNA and protein sequence of SUMO-Miniprotein2 construct:**

5’-
ATGGGTCATCACCATCATCATCACGGGTCGGACTCAGAAGTCAATCAAGAAGCCAAGCC
AGAGGTCAAGCCAGAAGTCAAGCCTGAGACTCACATCAATTTAAAGGTGTCCGATGGAT
CTTCAGAGATCTTCTTCAAGATCAAAAAGACCACTCCTTTAAGAAGGCTGATGGAAGCGT
TCGCTAAAAGACAGGGTAAGGAAATGGACTCCTTAAGATTCTTGTACGACGGTATTAGAA

TTCAAGCTGATCAGACCCCTGAAGATTTGGACATGGAGGATAACGATATTATTGAGGCTC
ACCGCGAACAGATTGGAGGTGCTAGCGCATGTGCTGAAGATTTAAAAGAGAGGCTAAAG
AAGCTGGGTTGTAGCGAAGAGTGCCGTCAGCGTTTGGAAAAAATGTGCAAAGAGGGCA
CCTGTGAGGACGCGGAACGTATGGCTCGCAACTGCGAGTCCTAA-3’

MGHHHHHHGSDSEVNQEAKPEVKPEVKPETHINLKVSDGSSEIFFKIKKTTPLRRLMEAFAK
RQGKEMDSLRFLYDGIRIQADQTPEDLDMEDNDIIEAHREQIGGASACAEDLKERLKKLGCS
EECRQRLEKMCKEGTCEDAERMARNCES*

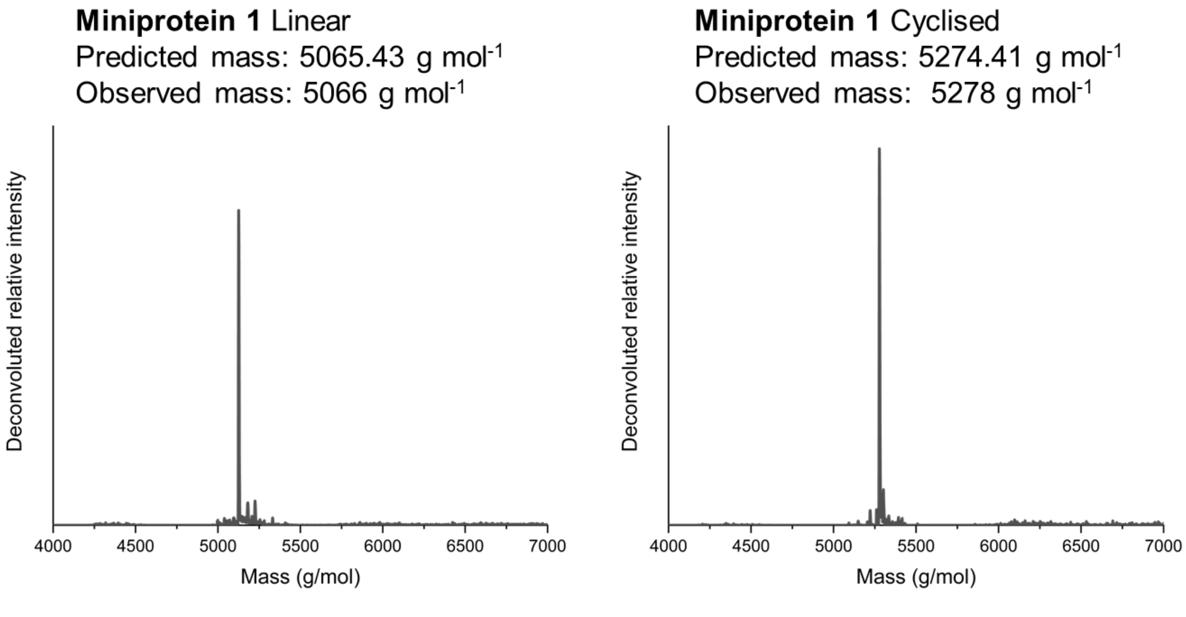


**Figure S1** – Deconvoluted ESI-MS spectra of **Miniprotein1** Linear and Cyclised.

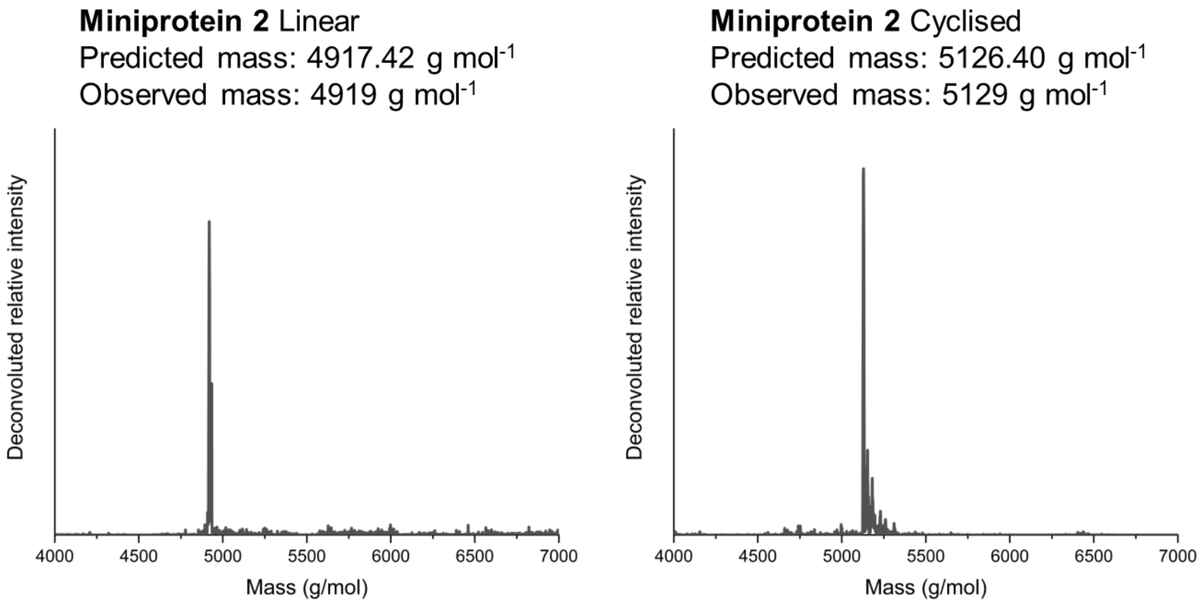


**Figure S2** – Deconvoluted ESI-MS spectra of **Miniprotein2** Linear and Cyclised.

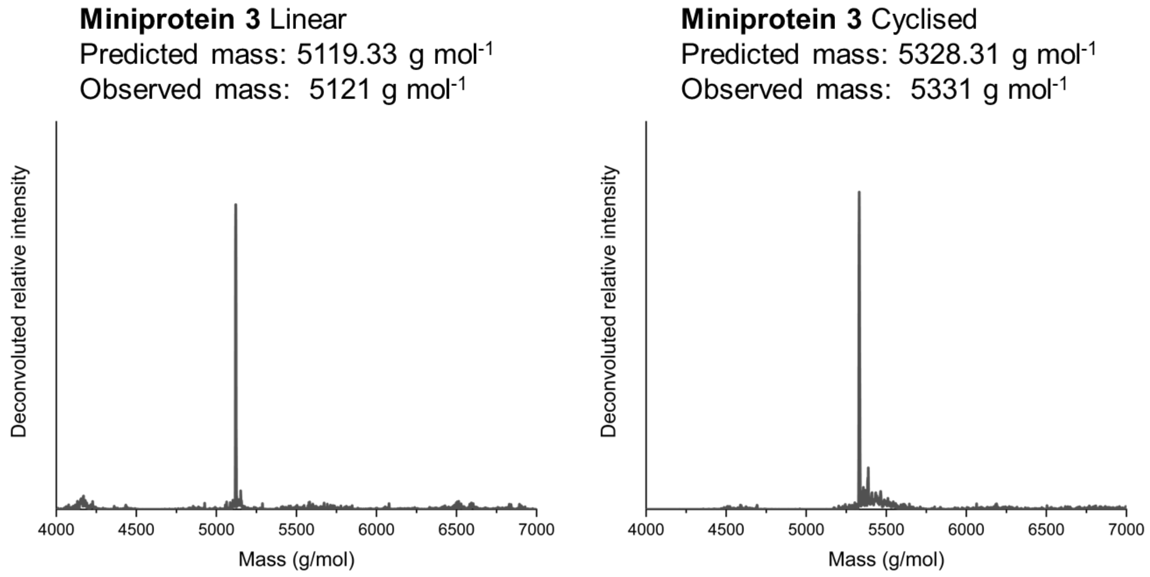


**Figure S3** – Deconvoluted ESI-MS spectra of **Miniprotein3** Linear and Cyclised.

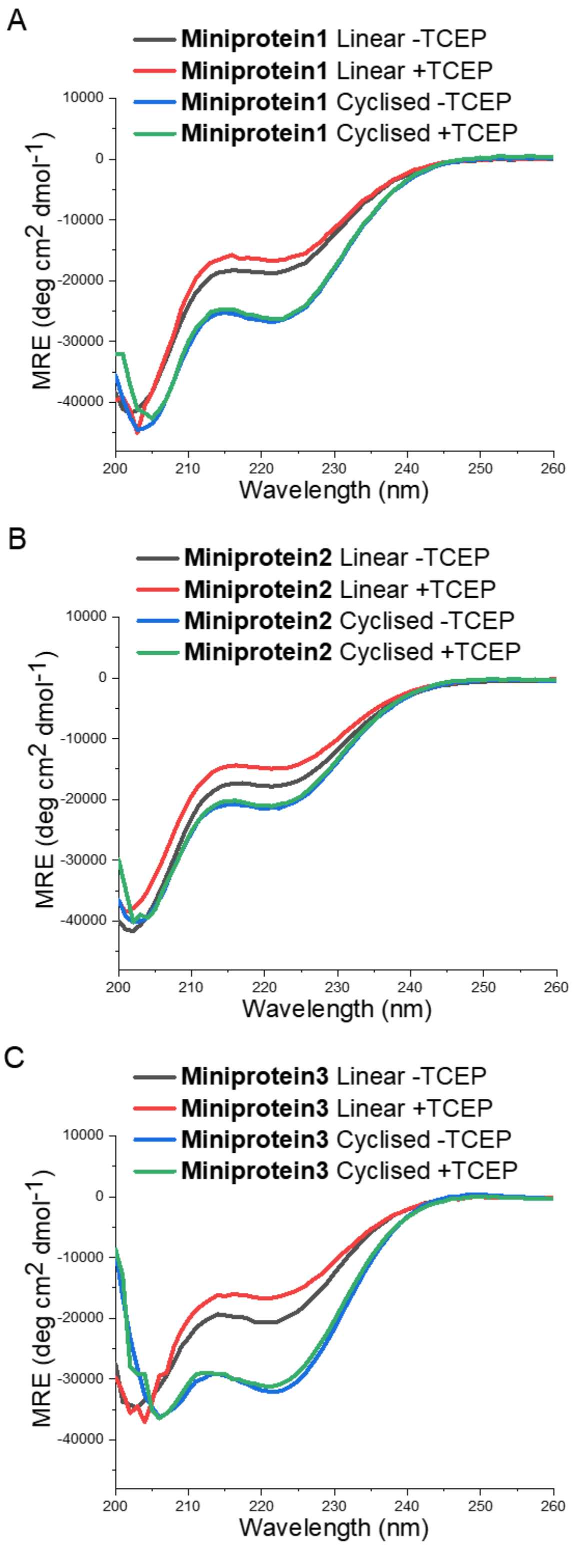


**Figure S4** – CD spectra of linear and cyclised miniproteins (100 μM) in the presence and absence of TCEP.

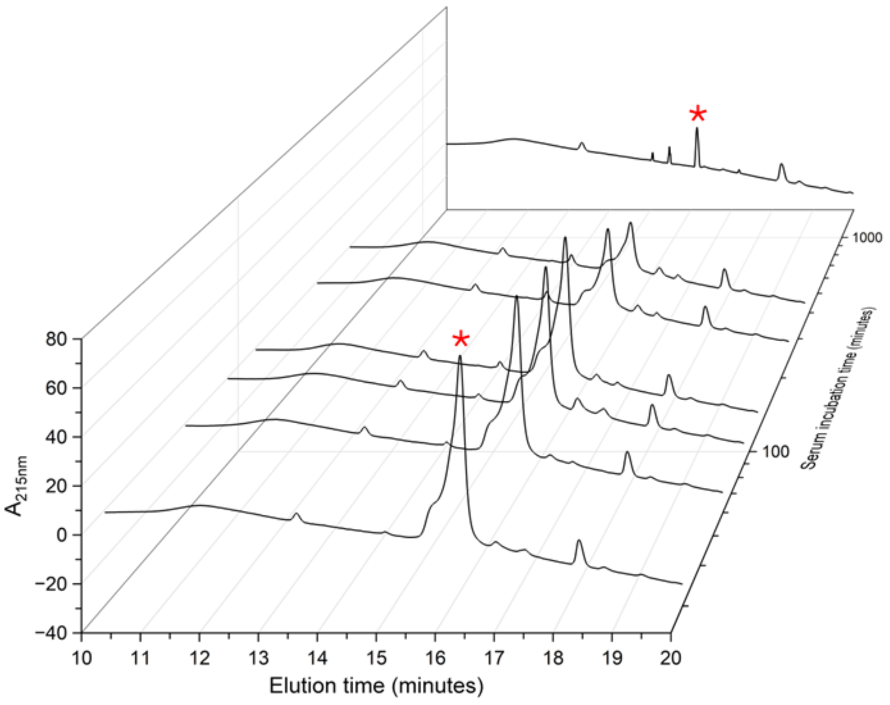


**Figure S5** – HPLC chromatogram of **Miniprotein3** Cyclised extracted after human serum incubation over the time course, as exemplar data for the generation of **Figure 4C**. Red star indicates peak corresponding to intact miniprotein, showing depletion over time.

A

**SUMO-Miniprotein 1** Linear
Predicted mass: 17366.44 g mol$^{-1}$
Observed mass: 17367 g mol$^{-1}$

**SUMO-Miniprotein 1** Cyclised
Predicted mass: 17575.42 g mol$^{-1}$
Observed mass: 17579 g mol$^{-1}$

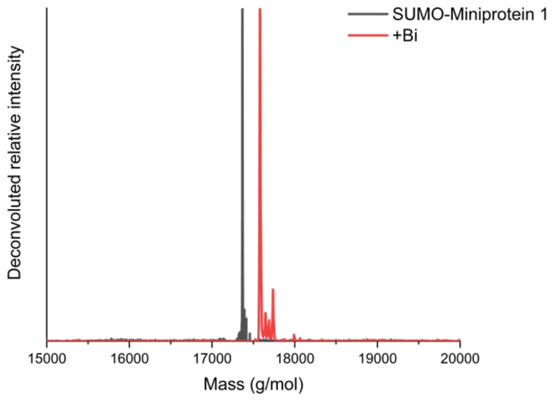


B

**SUMO-Miniprotein 3** Linear
Predicted mass: 17420.58 g mol$^{-1}$
Observed mass: 17423 g mol$^{-1}$

**SUMO-Miniprotein 3** Cyclised
Predicted mass: 17838.54 g mol$^{-1}$
Observed mass: 17840 g mol$^{-1}$

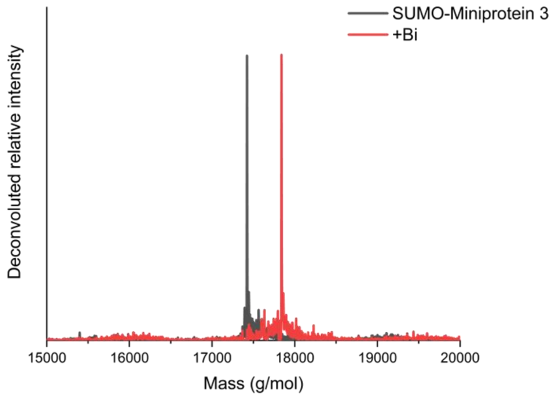


**Figure S6** – ESI-MS spectra showing intracellular bismuth cyclisation of (A) SUMO-Miniprotein1 and (B) SUMO-Miniprotein3. Proteins were recombinantly expressed in *E. coli* with or without 250 μM $BiK_3[citrate]_2$ supplementation, with ESI-MS indicating an increase in mass corresponding to the reaction with one or two bismuth atoms. N-terminal methionine truncation was observed upon recombinant expression.

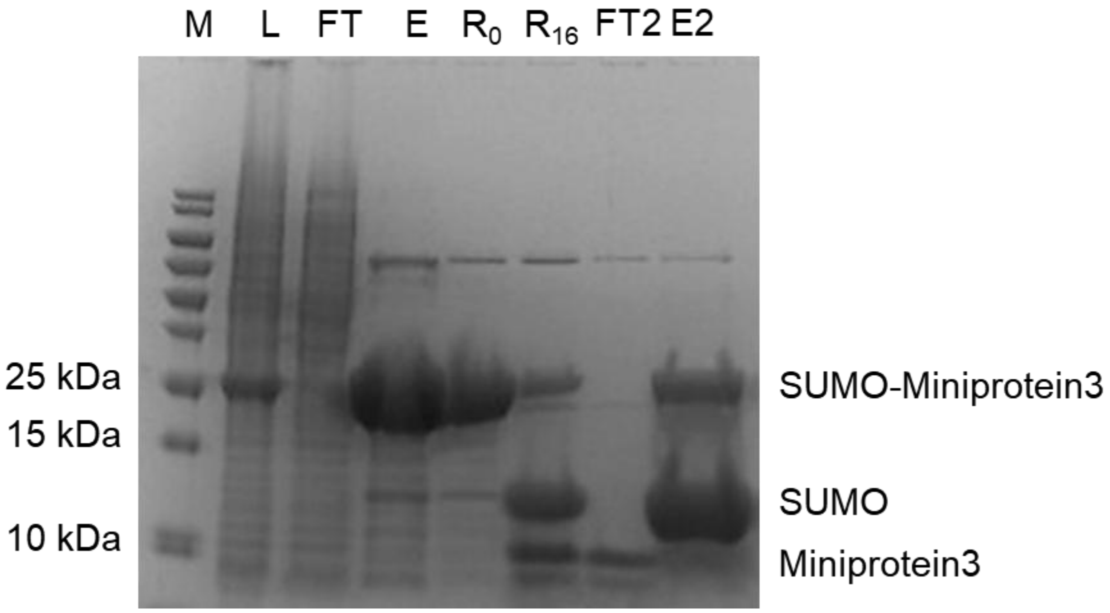


**Figure S7** – SDS-PAGE illustrating purification of **Miniprotein3** from recombinant expression. L: Lysate of cells expressing the SUMO-miniprotein3 fusion construct. FT: IMAC flow through showing the 6xHis-tagged construct bound to the column. E: SUMO-Miniprotein3 elution from column with imidazole. $R_0$: ULP1 cleavage reaction starting material after dilution of the IMAC column elution. $R_{16}$: ULP1 cleavage reaction after 16 hours showing starting material cleavage into constituent SUMO and Miniprotein3. FT2: Second IMAC column flow through, whereby the cleaved Miniprotein3 does not bind to the column, this sample underwent a final HPLC purification step. E2: Second IMAC column elution with imidazole, with unreacted fusion protein and cleaved SUMO contaminants removed. SUMO-Miniprotein3 and Miniprotien3 cleaved had a higher apparent molecular weight than the predicted mass however ESI-MS confirmed correct masses for these constructs. This likely indicates that Miniprotein3 Cyclised was not fully denatured and complexed with SDS due to its chemically constrained, stable folded structure.

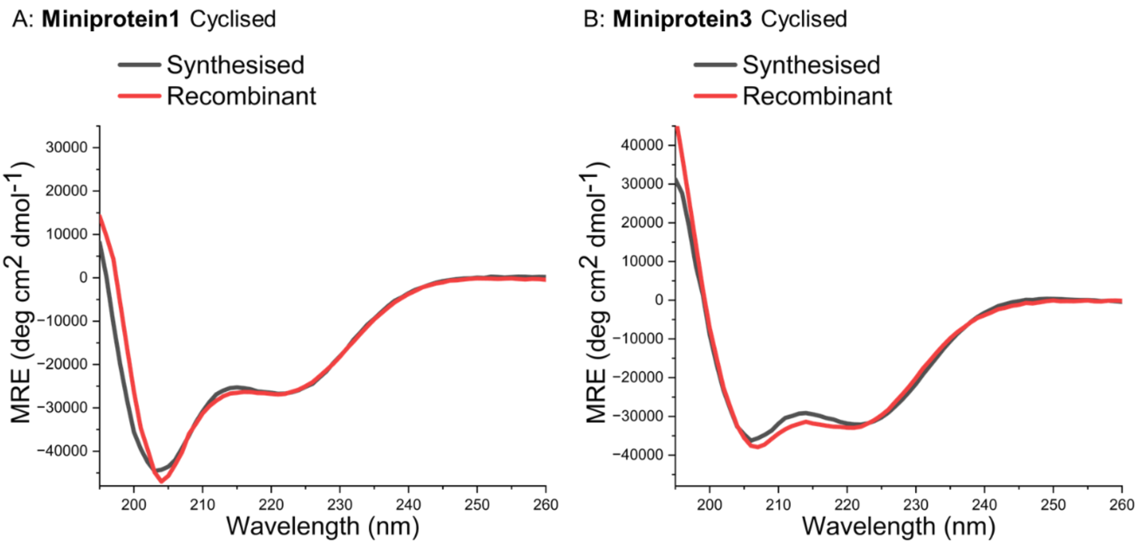


**Figure S8** – CD spectra of synthesised and recombinantly expressed (A) **Miniprotein1** Cyclised and (B) **Miniprotein3** Cyclised, illustrating that intracellular cyclisation produces highly similar secondary structure as synthesis.

| **$BiBr_3$ (µM)** | **Cyclic / linear ratio** |
|---|---|
| 25 | 0.9 |
| 50 | 1.3 |
| 75 | 3.0 |
| 100 | 1.4 |
| 250 | 14.7 |
| **$BiK_3$[citrate]$_2$** | |
| 50 | 2.5 |
| 100 | 8.4 |
| 250 | 21.0 |
| 500 | 27.9 |
| 1000 | 113.0 |

***Table S1 –*** *Cyclic-to-linear peptide ratios for recombinantly expressed Test Peptide following supplementation of bacterial growth media with bismuth salts, determined by LC-MS. Ratios were calculated from MS peak intensities and used as a relative measure of intracellular cyclisation efficiency. Increasing bismuth salt concentration resulted in higher cyclisation efficiency, with* $BiK_3$*[citrate]*$_2$ *exhibiting reduced toxicity and enabling higher ratios over a broader concentration range.*
